# Observation of an α-synuclein liquid droplet state and its maturation into Lewy body-like assemblies

**DOI:** 10.1101/2020.06.08.140798

**Authors:** Maarten C. Hardenberg, Tessa Sinnige, Sam Casford, Samuel Dada, Chetan Poudel, Lizzy Robinson, Monika Fuxreiter, Clemens Kaminksi, Gabriele S. Kaminski Schierle, Ellen A. A. Nollen, Christopher M. Dobson, Michele Vendruscolo

## Abstract

Misfolded α-synuclein is a major component of Lewy bodies, which are a hallmark of Parkinson’s disease. A large body of evidence shows that α-synuclein can self-assemble into amyloid fibrils, but the relationship between amyloid formation and Lewy body formation still remains unclear. Here we show, both *in vitro* and in a *C. elegans* model of Parkinson’s disease, that α-synuclein undergoes liquid-liquid phase separation by forming a liquid droplet state, which converts into an amyloid-rich hydrogel. This maturation process towards the amyloid state is delayed in the presence of model synaptic vesicles *in vitro*. Taken together, these results suggest that the formation of Lewy bodies is linked to the arrested maturation of α-synuclein condensates in the presence of lipids and other cellular components.

## Introduction

Parkinson’s disease (PD) is the most common neurodegenerative movement disorder, affecting over 2% of the world’s population over 65 years of age [1, 2]. The molecular origins of this disease are not fully understood, although they have been closely associated with dysregulation of the behaviour of α-synuclein [1, 3, 4], a disordered protein whose function appears to involve the regulation of synaptic vesicle trafficking [5-8]. This gap in our knowledge of the disease aetiology is no doubt partly responsible for the lack of disease modifying therapies for PD [9-12].

The pathological hallmark of PD is the presence of Lewy bodies within dopaminergic neurons in the brains of affected patients [13]. Although misfolded α-synuclein is a major constituent of Lewy bodies [14, 15], ultrastructural and proteome studies have indicated that these aberrant deposits contain a plethora of cellular components, including lipid membranes and organelle fragments [16-20]. These findings have led to the suggestion that the processes associated with the formation of Lewy bodies could be major drivers of neurotoxicity in PD [16, 18, 21].

Since it has also been shown that α-synuclein can aggregate into ordered amyloid fibrils *in vitro* [22-26] and *in vivo* [27-32], we asked how such α-synuclein aggregation can be reconciled with the formation of highly complex and heterogeneous assemblies such as Lewy bodies. We hypothesised that α-synuclein may be capable of irreversibly capturing cellular components through liquid-liquid phase separation, as this mechanism has been shown to drive the self-assembly of various disease-associated proteins on-pathway to the formation of solid aggregates [33-36]. Under healthy conditions, the condensation of proteins into a dense liquid droplet state through liquid-liquid phase separation is normally reversible, and exploited in a variety of ways to carry out cellular functions, including RNA metabolism, ribosome biogenesis, DNA damage response and signal transduction [33, 34, 37]. Upon dysregulation, however, liquid droplets can mature into gel-like deposits, which irreversibly sequester important cellular components and lead to pathological processes [38]. This phenomenon has been observed for FUS and TDP-43 in ALS [38, 39] and more recently also for tau in Alzheimer’s disease [40-42]. Our results show that α-synuclein can undergo a similar process by forming a dense liquid droplet state, which matures into a gel-like state rich in amyloid structure, consistently with very recent results [43].

## Results

### α-synuclein forms non-amyloid inclusions in *C. elegans*

To understand the initial events in the α-synuclein aggregation process, we assessed its self-association using confocal fluorescence lifetime imaging (FLIM) in the nematode worm *Caenorhabditis elegans*. The fluorescence lifetime of a fluorophore has been previously shown to be an accurate measure of protein self-assembly and amyloid formation, independent of protein concentration or fluorescence intensity [44-46]. Given the optical transparency and well-established genetics of *C. elegans*, we sought to compare the fluorescence lifetime of a human α-synuclein-YFP fusion protein expressed in body wall muscle cells (OW40 worm model, see **Methods**). In line with previous reports [47], we observed age-dependent coalescence of α-synuclein-YFP into distinct cytoplasmic inclusions (**Figure 1A**). Previous fluorescence recovery after photobleaching (FRAP) and FLIM studies have suggested that α-synuclein in inclusions exists in a mobile, non-amyloid state [44-47]. Here, we performed high resolution FLIM to be able to differentiate the protein localised at inclusions from the diffuse (non-aggregated) pool (**Figure 1A**). Our analysis revealed that the self-association state of α-synuclein-YFP in inclusions was similar to that of the diffuse state for the majority of the nematode’s adult life (days 1-11 of adulthood), indicating that α-synuclein assemblies are predominantly non-amyloid in *C. elegans* (**Figure 1B,C**). Only at old age, α-synuclein-YFP inclusions started to show increased amyloid-like features (days 13-15 of adulthood), an observation supported by the simultaneous appearance of proteinase K resistant protein species (**Figure 1D**).

**Figure 1.**
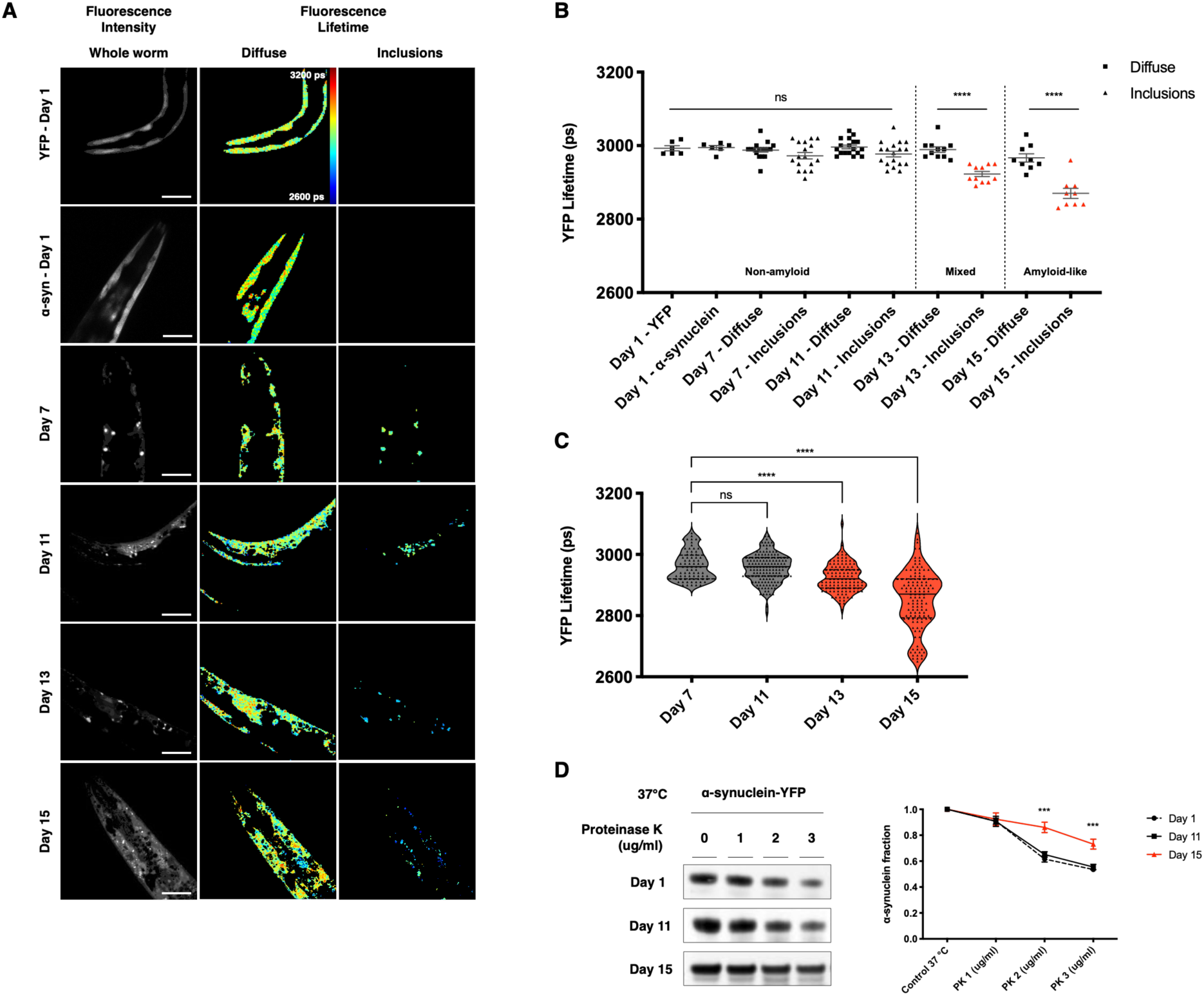
α-Synuclein forms non-amyloid inclusions in *C. elegans*. **(A)** Age-dependent coalescence of α-synuclein-YFP into cytoplasmic inclusions with distinct lifetime profiles between inclusions and diffuse signal. The scale bar represents 50 μm, rainbow scale bar 2600 (blue) - 3200 (red) picoseconds (ps). **(B)** Analysis of the high-resolution FLIM revealing that α-synuclein assemblies are predominantly non-amyloid in *C. elegans*. At day 13 and 15 a shift towards lifetimes associated with the amyloid-state can be observed. Each square and triangle represents one animal from the population. **(C)** Violin plots for all individual inclusions from the animals in (B). A distinct population of lower lifetime inclusions appears at old age. Black line = median. Dotted line = quartiles. **(D)** Appearance of proteinase K resistant protein species show α-synuclein-YFP with increased amyloid-like self-association in aged animals (days 13-15 of adulthood). α-Synuclein levels probed with the LB509 antibody (Methods) are shown (*left*). Quantification of protein levels corrected for loading control and normalised against untreated control (*right*). Results are mean ± SEM. One-way ANOVA. \*\*\**P* **≤** 0.001, ****P **≤** 0.0001, ns, not significant.

### Early α-synuclein inclusions in *C. elegans* have liquid-like properties

The presence of non-amyloid α-synuclein assemblies in *C. elegans* prompted us to further investigate their material properties. To this end, we used the aliphatic alcohol 1,6-hexanediol, which has recently been used as a tool to test the liquid properties of protein condensates *in vitro* as well as *in vivo* [48]. Hexanediol dissolves assemblies maintained by weak hydrophobic interactions, whereas solid-like amyloid structures remain intact. Exposure to 10% (w/v) hexanediol in *C. elegans* almost completely dissolved α-synuclein inclusions between days 1 and 11 of adulthood, reflecting the role of low-affinity interactions in inclusion assembly (**Figure 2A,B** and **Movie S1**). As before, inclusions in aged nematodes (days 13-15 of adulthood) appeared to be solid-like, as evidenced by decreased hexanediol-mediated dissipation. We then assessed whether α-synuclein assemblies in younger nematodes could reappear after hexanediol was washed out. Indeed, newly formed small inclusions were observed in washed worms after one day of recovery, reflecting the dynamic and liquid-like properties of α-synuclein assemblies in *C. elegans* (**Figure 2C,D**).

**Figure 2.**
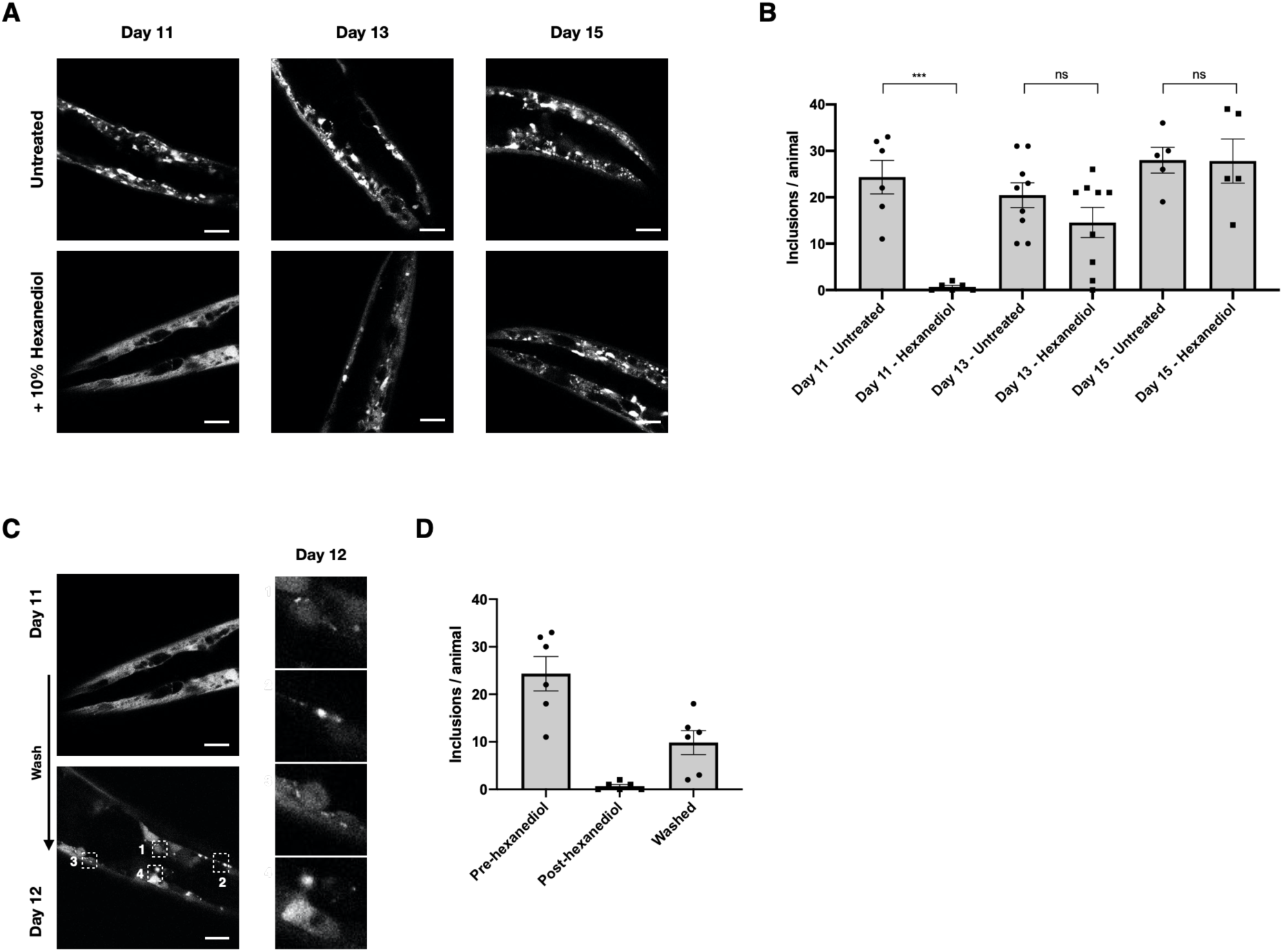
α-Synuclein inclusions in *C. elegans* have liquid-like properties. **(A)** Exposure to 10% (w/v) 1,6-hexanediol dissolves α-synuclein inclusions between days 1 and 11 of adulthood, reflecting the dependence of inclusion assembly on weak hydrophobic interactions. By contrast, inclusions in aged nematodes (days 13-15 of adulthood) appear to be solid-like, as evidenced by decreased hexanediol efficacy. **(B)** Quantification of images in (A). Each square or circle represents one animal from the population. (**C**) α-Synuclein assemblies in younger nematodes reappear after hexanediol is washed out. Newly formed small inclusions were observed in washed worms after one day of recovery, reflecting the dynamic and liquid-like nature of α-synuclein assemblies in *C. elegans.* (**D**) Quantification of images in (C). The scale bars represent 25 μm. Results are mean ± SEM. One-way ANOVA. \*\*\**P* **≤** 0.001, ****P **≤** 0.0001, ns, not significant.

### Liquid-liquid phase separation drives α-synuclein droplet formation *in vitro*

The results presented above indicate that α-synuclein is a component of liquid-like assemblies in *C. elegans*. To investigate whether this behaviour is mediated by the intrinsic biophysical properties of α-synuclein, we tested its capability of forming such assemblies *in vitro*. We incubated Alexa-488 labelled human α-synuclein at physiological pH and in the presence of polyethylene glycol (PEG), a commonly used crowding agent (**Methods**). When deposited on a glass surface, α-synuclein formed micron-sized droplets within several minutes (**Figures 3A** and **S1**, and **Movie S2**). Droplet formation was sensitive to ionic strength (**Figure S2**), which is likely the result of the involvement of the acidic C-terminal region of α-synuclein. This region is predicted to spontaneously phase separate via disordered interactions (**Figure S3**). To characterise whether α-synuclein droplets had properties of a liquid phase, we assessed two key biophysical features. First, we observed that two droplets readily fuse and relax into a larger droplet once they come in close proximity (<1 μm) (**Figure 3B** and **Movie S3**). Second, fluorescence recovery after photobleaching (FRAP) of a small area within the droplet was followed by rapid recovery (t_1/2_ = 5 s), reflecting local rearrangement of α-synuclein molecules within the condensate (**Figure 3C** and **Movie S4**).

**Figure 3.**
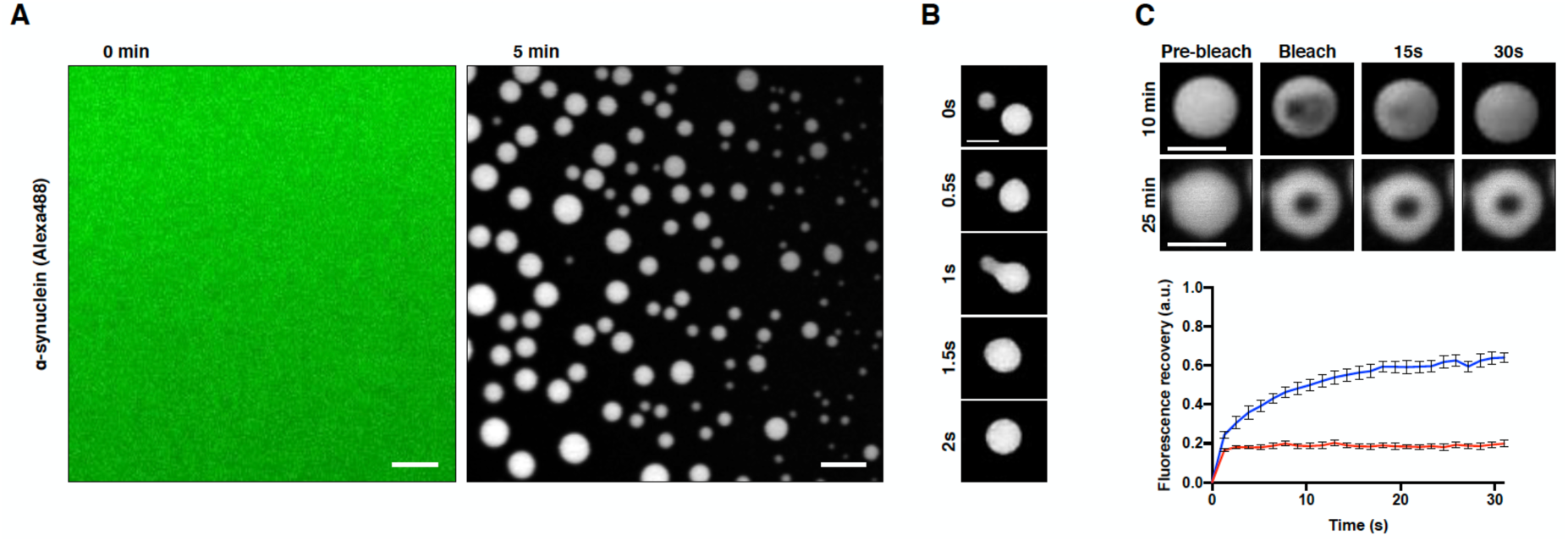
α-Synuclein undergoes liquid-liquid phase separation *in vitro*. **(A)** Wild-type human α-synuclein, supplemented with 1 mol% Alexa-488 labelled α-synuclein, forms micrometre sized droplets under physiological conditions. The scale bar represents 5 μm. **(B)** Fusion of two α-synuclein droplets in close proximity (<1 μm). The scale bar represents 1 μm. **(C)** Fluorescence recovery after photobleaching (FRAP) of a small area within the droplet. Red line represents aged droplets. The scale bar represents 1 μm; error bars represent SEM.

### The conversion of α-synuclein droplets into gel-like assemblies is associated with amyloid formation

The ability of α-synuclein molecules to rearrange within droplets declines over time (**Figure 3C**), which is in line with the thermodynamic drive of dense protein solutions to mature into more solid-like states, such as highly ordered amyloid fibrils [38, 41, 49, 50]. To test whether the maturation of α-synuclein droplets is associated with amyloid formation, we assessed this process using turbidity measurements in the presence of the amyloid-binding dye thioflavin T (ThT) (**Figure 4A,B**). Droplet formation as seen from increased turbidity at 540 nm was followed by increased ThT fluorescence, indicating that droplet ageing involves the formation of amyloid species. Furthermore, a hydrogel with amyloid-like features could be recovered at the end of each incubation period (**Figure 4C**). These results indicate that under physiological conditions, α-synuclein can undergo a phase transition into a dynamic liquid droplet state which converts into gel-like aggregates over time.

**Figure 4.**
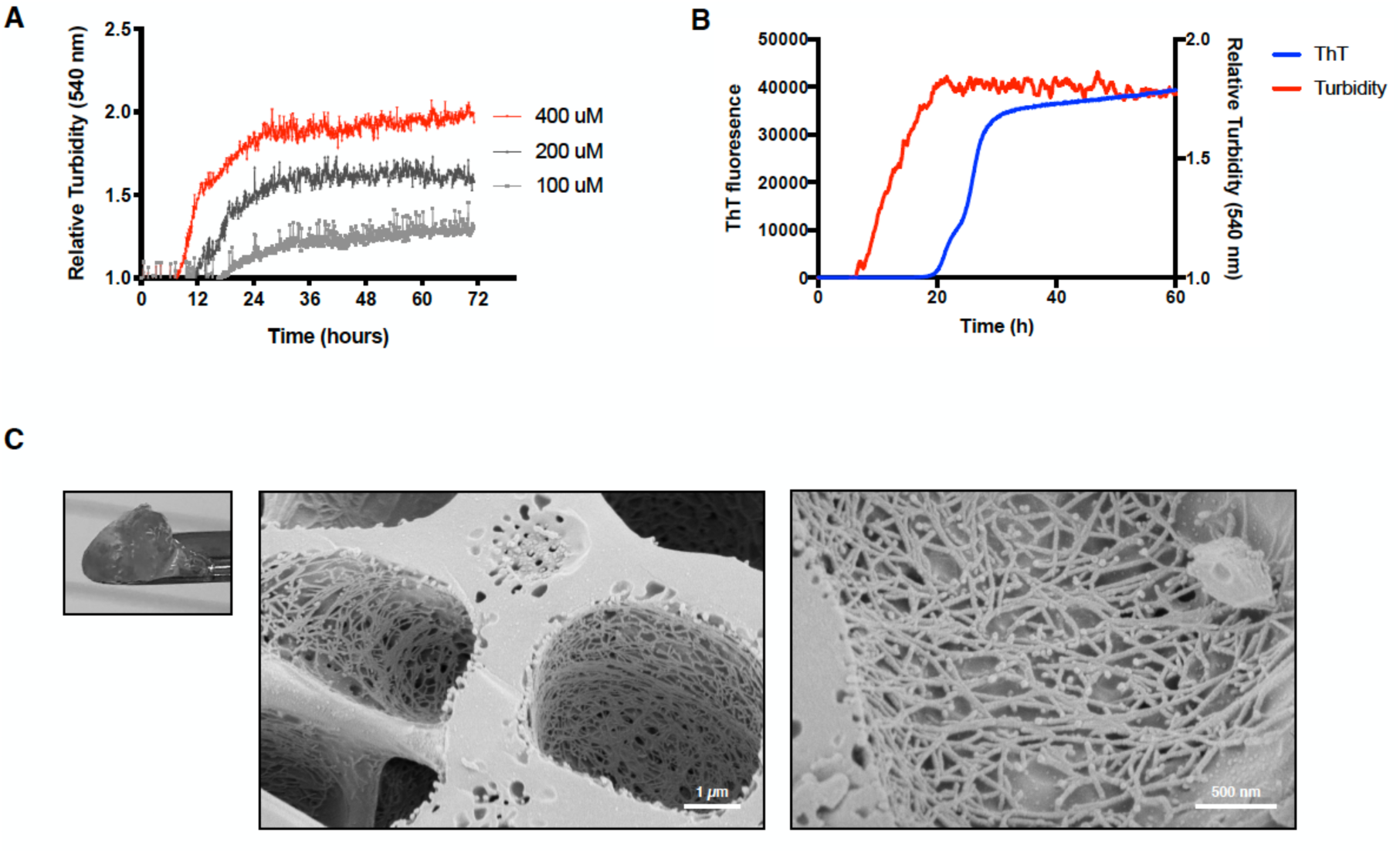
α-Synuclein droplets mature into amyloid-like hydrogels under *in vitro*. **(A)** α-Synuclein in the presence of 10% PEG becomes turbid, representing the formation of light-scattering objects such as droplets. Data points represent 5 min interval measurements, preceded by brief shaking. **(B)** α-Synuclein becomes positive for ThT following an increase in turbidity, indicating that the assemblies adopt an amyloid-like nature. **(C)** Scanning electron microscopy (cryo-SEM) image of α-synuclein hydrogel, formed at the end of the experiment shown in (B). Left panel shows the hydrogel. The morphology as seen by cryo-SEM (middle and right panels) indicates that hydrogels are rich in fibrillar structures. The scale bars represent 1 μm (*left)* and 500 nm *(right).*

### α-Synuclein droplets age more slowly when merged with liposomes

In dopaminergic neurons, α-synuclein is highly concentrated at pre-synaptic terminals where it can transiently bind a variety of lipid surfaces, including those of synaptic vesicles [5, 7, 8, 51]. These observations lead to the question how the presence of synaptic vesicles modulates the phase behaviour of α-synuclein. We thus incubated α-synuclein with liposomes with a composition mimicking that of synaptic vesicles [52] (**Methods**). To enable detection by fluorescence microscopy, liposomes were supplemented with a fluorescently labelled lipid, 1,2-dioleoyl-sn-glycero-3-phosphoethanolamine-N-(cyanine 5) (Cy5-DOPE) (**Methods**). Formation of α-synuclein droplets coincided with the appearance of condensates that were positive for the labelled lipid (**Figure 5A** and **Movie S5**), whereas liposomes alone either remained diffuse or formed irregular aggregates (**Figure 5B**). 3D rendering of the droplets showed that lipids are directly recruited into α-synuclein droplets (**Figure 5C** and **Movie S6**) which continue to behave as a liquid phase as evidenced by rapid fusion of two α-synuclein/lipid droplets (**Figure 5D**). Crucially, α-synuclein/lipid droplets appeared to be more resistant to ageing, as they showed a slower decline in FRAP when compared to α-synuclein droplets alone (**Figure 5E**). These results suggest that liposomes mimicking synaptic vesicles stabilise the liquid state of α-synuclein, which is expected as binding of α-synuclein to these membranes does not readily trigger amyloid formation *in vitro*, although the binding to other membranes could have an opposite effect [53].

**Figure 5.**
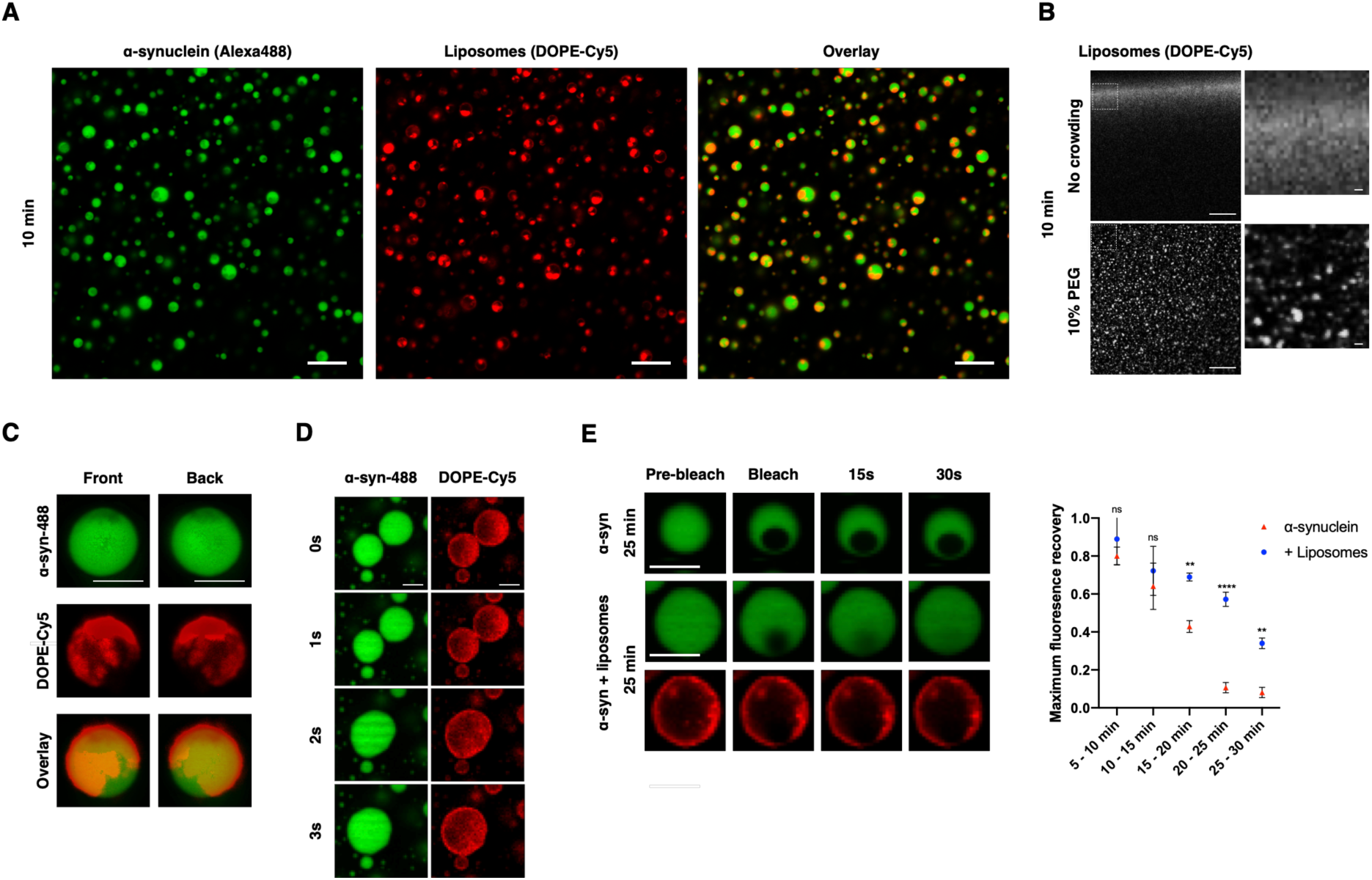
α-Synuclein droplets co-localise with liposomes. **(A)** α-Synuclein droplets co-localise with liposomes resembling the lipid composition of synaptic vesicles (DOPE:DOPC:DOPS) labelled with DOPE-Cy5. The scale bar represents 5 μm. **(B)** Liposomes in buffer remain diffuse (*top*) or form aggregates (*bottom*). The scale bar represents 10 μm. **(C)** 3D rendering of the droplets showing that lipids are directly recruited into α-synuclein droplets. The scale bar represents 1 μm. **(D)** Fusion of two α-synuclein/liposome droplets in close proximity. The scale bar represents 1 μm. (**E**) α-synuclein/liposome droplets mature slower than droplets containing only α-synuclein. Each data point represents the average recovery out of three droplets for that timeframe. The scale bar represents 1 μm; error bars represent SEM. One-way ANOVA. \*\**P* **≤** 0.01, ****P **≤** 0.0001, ns, not significant.

## Discussion and Conclusions

We have shown that α-synuclein forms liquid-like condensates *in vivo* in a *C. elegans* model of Parkinson’s disease and *in vitro* at physiological concentrations and pH. Our FLIM analysis indicated that the inclusions formed by α-synuclein in *C. elegans* body wall muscle cells are largely non-amyloid and can be dissociated by 1,6-hexanediol. α-Synuclein droplets behave as a liquid phase *in vitro*, as individual α-synuclein molecules rapidly rearrange within the droplets and individual droplets were observed to fuse and relax back into a spherical shape.

These observations can be linked to the increasing evidence that the phenomenon of liquid-liquid phase separation underlies the organisation of proteins at the synapse [54-56]. It has also been shown that α-synuclein clusters synaptic vesicles and regulates the synaptic vesicle pool [5-8]. In this context, our results suggest that droplet formation by α-synuclein may be involved in a physiological mechanism to cluster synaptic vesicles, possibly in conjunction with other proteins such as synapsins, which have also been shown to phase-separate into liquid-like assemblies [55]. Indeed, synapsins are implicated in synaptic function in conjunction with synuclein [57, 58] and are components of Lewy bodies [59].

We have also shown that the droplet state can undergo a maturation process into a gel-like state rich in amyloid structure, which is reminiscent of the pathological state seen in Lewy body pathologies [14, 15, 60]. This phenomenon is expected on the grounds that the transition from the droplet state to the amyloid state takes place through a maturation process, known as Ostwald ripening [61, 62], and it is consistent with the recent report that α-synuclein can form hydrogels [63, 64]. From a thermodynamic point of view, it is important to note that the gel-like assembly is not a stable state in addition to the native, the droplet and the amyloid states, but a slowly evolving conformation on pathway to the amyloid state (**Figure 6A**). This ageing process can become very slow depending on the complexity of the composition, and one can speculate that Lewy bodies are in a condition of nearly arrested maturation. In this context, we found that the sequestration of liposomes mimicking synaptic vesicles slows the maturation process, which is consistent with the observation that various lipid vesicles and membranes are found in Lewy bodies [16, 17]. We can thus suggest that the effects of such membranous structures on the ageing process of the droplets involve a slowing down of the amyloid conversion of the gel-like α-synuclein assemblies. This process could create a metastable toxic state, as the gel-like assemblies can be a reservoir for cytotoxic α-synuclein oligomers [64].

**Figure 6.**
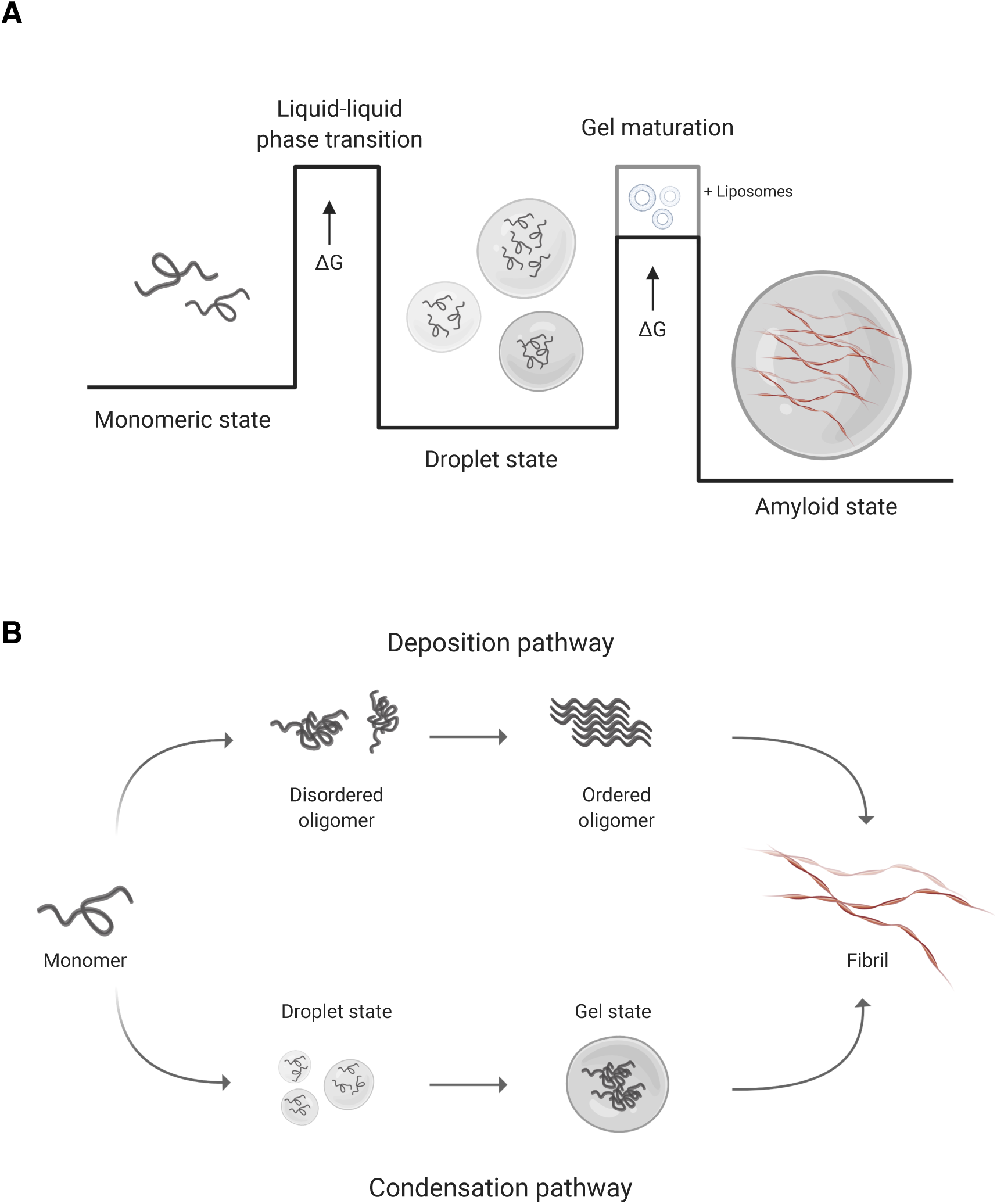
Schematic illustration of the thermodynamics and kinetics of α-synuclein aggregation. **(A)** α-Synuclein can populate the monomeric state, the droplet state and the amyloid state. The amyloid state is likely to be the thermodynamically most stable state under cellular conditions, but its kinetic accessibility is reduced by the presence of free energy barriers between the native and droplet states, and by the droplet and amyloid states. The latter free energy barrier can be crossed through a maturation process that involves the formation of gel-like amyloid-rich assemblies that gradually age into the amyloid state. This slow conversion can become altogether arrested in the presence of cellular components, possibly resulting in the formation of Lewy bodies. **(B)** The conversion of α-synuclein from the monomeric to the amyloid state can proceed through two distinct pathways. In the ‘deposition pathway’, α-synuclein forms initially small disordered oligomers that then convert into ordered oligomers, which then grow into amyloid fibrils. In the ‘condensation pathway’ α-synuclein forms first a droplet state, which then gradually mature into the amyloid state going through a gel intermediates. Although both pathways can be homogeneous, with α-synuclein as the sole component of the system, in complex environment they are most likely heterogeneous, with other cellular components taking part in the process.

We also note that the conversion of α-synuclein between the native state and the amyloid state can take place directly through a ‘deposition pathway’ (**Figure 6B**) following a more conventional nucleation-growth process [65], where the nucleation step can be catalysed by the presence of charged lipid membranes [66], rather than the ‘condensation pathway’ described here (**Figure 6B**). Whether α-synuclein follows primarily the deposition route to the amyloid state or the condensation route through the droplet state is likely determined by the environmental conditions. One such condition could involve the lipid composition of membranes, since this determines the rate of α-synuclein nucleation, with membranes mimicking synaptic vesicles being unable to drive nucleation [53]. A decline in synaptic vesicles density, as observed for some PD mutations through altered synaptic protein function [67], could therefore shift α-synuclein membrane binding with adverse consequences.

In conclusion, we have shown that α-synuclein undergoes liquid-liquid phase separation to adopt a droplet state and that this process may lead to the formation of aberrant gel-like assemblies with features of Lewy bodies.

## Materials and Methods

### *C. elegans* strains and maintenance

Standard conditions were used for the propagation of *C. elegans* [68]. We used the OW40 (zgIs15[unc-54p::α-synuclein::YFP] IV) strain in which α-synuclein is fused to yellow fluorescent protein (YFP) [47]. OW40 animals were synchronised by hypochlorite bleaching, hatched overnight in M9 buffer (3 g/L KH_2_PO_4_, 6 g/L Na_2_HPO_4_, 5 g/L NaCl, 1 mM MgSO_4_) and subsequently cultured at 20 °C on nematode growth medium (NGM) plates (1 mM CaCl_2_, 1 mM MgSO_4_, 5 μg/mL cholesterol, 250 mM KH_2_PO_4_ [pH 6], 17 g/L agar, 3 g/L NaCl, 7.5 g/L casein), which were seeded with the *Escherichia coli* strain OP50. At L4 stage, larvae were placed on nematode growth medium (NGM) plates containing 5-fluoro-2’deoxy-uridine (FUDR, Sigma) (75 μM) to inhibit the growth of offspring.

### Fluorescence lifetime imaging (FLIM)

At indicated timepoints, transgenic worms were washed off NGM plates in M9 buffer and mounted on 2.5% agarose pads on glass microscopy slides and immobilised using 10 mM levamisole hydrochloride (Merck). Slides were then mounted onto a modified, confocal-based platform (Olympus FV300-IX700) integrated with a time-correlated single photon counting (TCSPC) module (SPC-830, B&H). Images of the anterior region of the worm were acquired at 60X (PLAPON 60XOSC2, 1.4NA, Olympus) magnification. For excitation of YFP, the output of a pulsed supercontinuum source (WL-SC-400-15, Fianium Ltd.) operating at 40 MHz repetition rate and filtered using a FF03-510/20 (Semrock Inc.) bandpass filter was used. YFP fluorescence emission was filtered with a FF01-542/27 (Semrock Inc.) bandpass filter. Photons were acquired for two minutes to create a single 256×256 image with 256 time bins. Inclusions were detected using a custom-designed image-processing pipeline in ImageJ (NIH). Briefly, cytoplasmic background was reduced through application of a Gaussian convolution filter and binarized according to the Chow and Kaneko adaptive thresholding method [69]. Subsequently, clustered inclusions were separated using a classic watershed segmentation algorithm. The resulting image was then used to create a mask of the signal representing inclusions and subtracted from the total signal, creating a separate mask for the diffuse signal. Lifetime analysis of the processed images was carried out using FLIMFit (Imperial College London Photonics). FLIM data were fitted with a single exponential decay function to extract the fluorescence lifetime in each pixel using FLIMFit v4.12 [70].

### Proteinase K assay

For proteinase K digestion assays, worms were lysed using a Balch homogeniser in NP-40 lysis buffer (150 mM NaCl, 1.0% NP-40, 50 mM Tris pH 8.0), as described previously [71]. Lysates were kept on ice before being exposed to increasing proteinase K concentrations (0, 1, 2, 3 μg/μl) for 30 minutes at 37 °C. Following digestion, samples were immediately heated at 100 °C for 5 minutes in Laemmli buffer, separated on NuPAGE Novex 4-12 % Bis-Tris Protein Gels (Life Technologies) and transferred to nitrocellulose membranes using an iBlot Dry Blotting System (Life Technologies). To improve α-synuclein immunodetection, the membrane was treated with phosphate-buffered saline (PBS) containing 0.4% paraformaldehyde for 30 min at room temperature, followed by blocking for 1 h with 5% skim milk in PBS. Membranes were probed with the LB509 anti-α-synuclein antibody (1:1000; Abcam) overnight at 4 °C and visualised with a fluorescent secondary antibody (goat-anti-mouse IgG Alexa 488-conjugated (1:5000, A11029, Life Technologies)). Fluorescent bands were detected on a Typhoon FLA 7000 (GE Life Sciences) imaging system. Images were quantified in ImageJ (NIH).

### Hexanediol experiments

To achieve slow diffusion of 1,6-hexanediol into the worms without eliciting rapid toxic effects, animals were washed off NGM plates in M9 buffer and embedded in 20 μL of a thermo-reversible hydrogel (CyGEL™, Biostatus) supplemented with 10% (w/v) 1,6-hexanediol (Sigma) and 10 mM levamisole hydrochloride (Merck) on a glass-bottom imaging dish (MatTek P35G-1.5-14-C). For control experiments, worms were embedded in hydrogel with 10% dH_2_O and 10 mM levamisole hydrochloride. Glass-bottom dishes were subsequently sealed with a 12 mm x 12 mm? 1.5 cover glass (Fisherbrand) and left at room temperature for 1 hour. After incubation, worms were imaged on a Leica SP8 upright confocal microscope using a 40x/1.3 HC PL Apo CS oil objective (Leica Microsystems). For recovery experiments, imaging dishes were cooled on ice until the hydrogel liquified and immediately diluted in M9 buffer. Worms were then washed 2x in M9 buffer, before being left to recover on seeded NGM plates for 24 hours. Hereafter, worms were imaged as described above.

### α-synuclein purification and labelling

Wild type and A90C α-synuclein were purified from *E. coli* expressing plasmid pT7-7 encoding for the protein as previously described [23, 72]. Following purification, the protein was concentrated using Amicon Ultra-15 Centrifugal Filter Units (Merck Millipore). A90C α-synuclein was labelled with 1.5-fold molar excess Alexa Fluor 488 C_5_ maleimide (Life Technologies) overnight at 4 °C. The excess dye was removed on a Sephadex G-25 desalting column (Sigma) as described previously [23], and buffer exchanged into 25 mM Tris-HCl (pH 7.4).

### Liquid-liquid phase separation assay

To induce droplet formation, non-labelled wild type α-synuclein was mixed with Alexa Fluor 488 labelled A90C α-synuclein at a 10:1 molar ratio in 25 mM Tris-HCl (pH 7.4), 50 mM NaCl, 1 mM DTT and 10% polyethylene glycol (PEG) (Thermo Fisher Scientific) at 20 °C, unless indicated otherwise. The final mixture was pipetted on a 35 mm glass-bottom dish (P35G-1.5-20-C, MatTek Life Sciences) and immediately imaged on a Leica TCS SP5 confocal microscope using a 40x/1.3 HC PL Apo CS oil objective (Leica Microsystems). The excitation wavelength was 488 nM for all experiments. All images were processed and analysed in ImageJ (NIH).

### Fluorescence recovery after photobleaching

Fluorescence recovery after photobleaching (FRAP) was performed on the setup described above for the liquid-liquid phase separation assay, under the same experimental conditions. Bleaching was done using the 488 nm laser at 50% intensity, to obtain ±50-60% relative photobleaching. Images were captured at 600 ms intervals, following a 1.8 s (3 frame) pre-bleach sequence and a 1.2 s (2 frame) spot bleach covering 30-50% of the droplet area. Intensity traces of the bleached area were background corrected and normalised.

### Liposome preparation

Small unilamellar vesicles (SUVs) were prepared to approximate the lipid composition of synaptic vesicles [52], 50 mol% DOPC (1,2-dioleoyl-sn-glycero-3-phosphocholine), 30 mol% DOPE (1,2-dioleoyl-sn-glycero-3-phosphatidylethanolamine), 20 mol% DOPS (1,2-dioleoyl-sn-glycero-3-phospho-L-serine), and supplemented with 1 mol% DOPE-Cy5 (1,2-dioleoyl-sn-glycero-3-phosphoethanolamine-N-(Cyanine 5)) (Avanti Polar Lipids). Lipids were mixed in chloroform and dried under a mild stream of nitrogen gas. Dried lipids were then lyophilised for 3 h (VWR, Avantor), before being rehydrated in 25 mM Tris-HCl (pH 7.4). Liposomes were formed by ten consecutive freeze-thaw cycles (−196 °C to 30°C). To form uniformly sized liposomes, the mixture was extruded 20 times through 50 nm-diameter polycarbonate filters (Avanti Polar Lipids). Liposome size distribution was confirmed using dynamic light scattering (Average size ± 50 nm) and re-extruded when standard deviation > 20 nm.

### Turbidity and thioflavin T (ThT) assay

Wild type α-synuclein was mixed with 20 μM ThT in 25 mM Tris-HCl (pH 7.4), 50 mM NaCl, and 10% PEG (Thermo Fisher Scientific) at 20 °C. All samples were prepared in low binding test tubes (Eppendorf), after which each sample was pipetted in triplicate into a 96-well half-area, low-binding, clear bottom plate (Corning) on ice?. Assays were initiated by placing the 96-well plate at 20 °C under intermediate (5 min) shaking (100 rpm, 10 s) conditions in a plate reader (Fluostar Omega or Fluostar Optima, BMG Labtech). The absorbance at 540 nm was measured every 5 min over 72 h. ThT fluorescence was measured through the bottom of the plate with a 440 nm excitation filter and a 480 nm emission filter.

### Electron microscopy (cryo-SEM)

Cryo-SEM was performed on a Verios 460 scanning electron microscope (FEI/Thermo Fisher) equipped with a Quorum PP3010T cryo-transfer system. Hydrogel samples were pipetted into shuttle-mounted universal cryo-stubs and flash-frozen in slushed nitrogen. After transfer into the prep-chamber, samples were fractured, sublimed at -90 °C for 2 minutes, sputter-coated with a thin layer of platinum and transferred to the SEM cryo-stage. Both prep-chamber and cryo-stage were set to -140 °C. SE-imaging was performed at 1 keV accelerating voltage and 25 pA probe current using the Through-Lens-Detector (TLD) in immersion mode. Images were acquired with a pixel resolution of 1536 x 1024 pixels using a 300 ns dwell time/32 image integrations and using drift correction.

### Droplet predictions

The binding modes of α-synuclein, as the probability to undergo disorder-to-order (p_DO_) or disorder-to-disorder (p_DD_) transitions upon interactions, were predicted from the amino acid sequence using the FuzPred program (protdyn-fuzpred.org) [73]. The FuzPred-Droplet method was used to estimate the probability for spontaneous liquid-liquid phase separation based on disorder in the free and bound states.

### Statistical analysis

All statistical analysis was performed in GraphPad Prism 8 (GraphPad Software). Data are presented as means ± SEM from at least 3 independent biological replicates, unless indicated otherwise. Statistical significance between experimental groups was analysed either by 2-tailed Student’s *t* test or one-way ANOVA followed by Bonferroni’s multiple-comparison

## Acknowledgements

We would like to acknowledge Nicola Lawrence at the Gurdon Institute Imaging Facility for confocal microscopy support and Karen Müller at the Cambridge Advanced Imaging Centre for her assistance with cryo-SEM experiments. We wish to thank Wing Man for his help with the preparation of liposomes. G.S.K.S. acknowledges funding from the Wellcome Trust (065807/Z/01/Z) (203249/Z/16/Z), the UK Medical Research Council (MRC) (MR/K02292X/1), Alzheimer Research UK (ARUK) (ARUK-PG013-14), Michael J Fox Foundation (16238) and from Infinitus China Ltd.

## SUPPLEMENTARY INFORMATION

**Figure S1.**
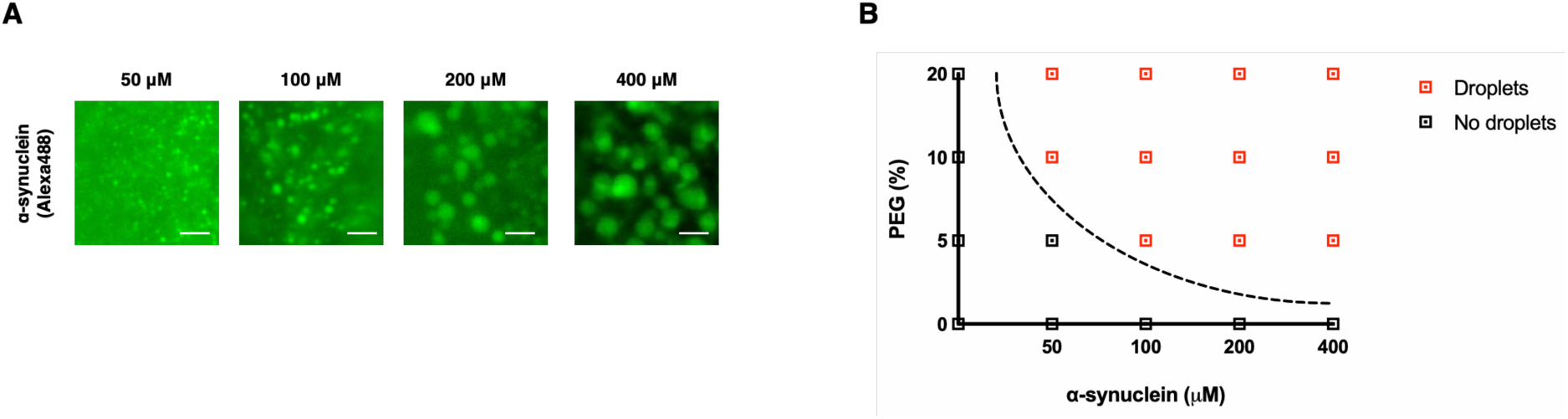
α-Synuclein forms droplets at various concentrations under crowding conditions. **(A)** Wide field microscopy images of α-synuclein forming micrometre-sized droplets in the presence of 10% PEG after 5 minutes incubation. Droplet size scales with protein concentration. The scale bar represents 5 μm. **(B)** Phase diagram of the droplet state as a function of α-synuclein and PEG concentration at 5 min post incubation.

**Figure S2.**
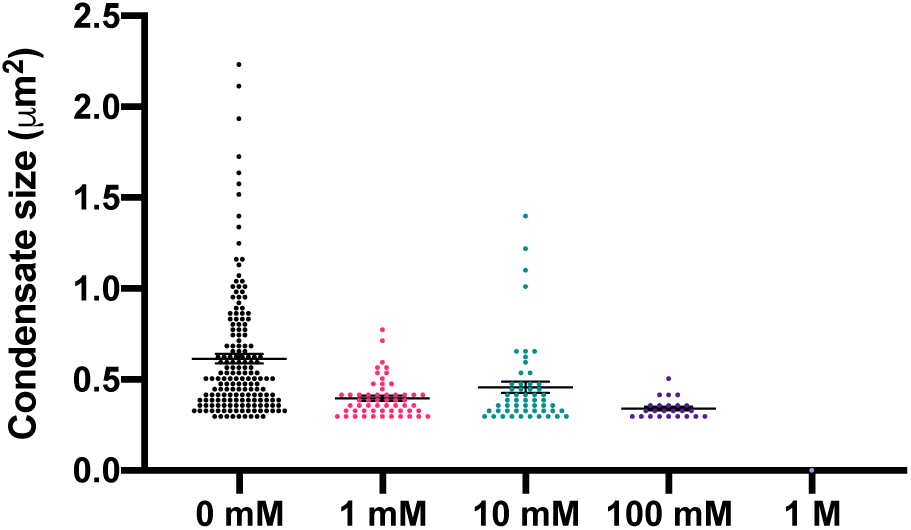
α-Synuclein droplet formation is sensitive to solution ionic strength. Dose-dependent distribution graphs representing the quantification of the effects of NaCl on the ?? of the average size of condensates at a 5 min time point.

**Figure S3.**
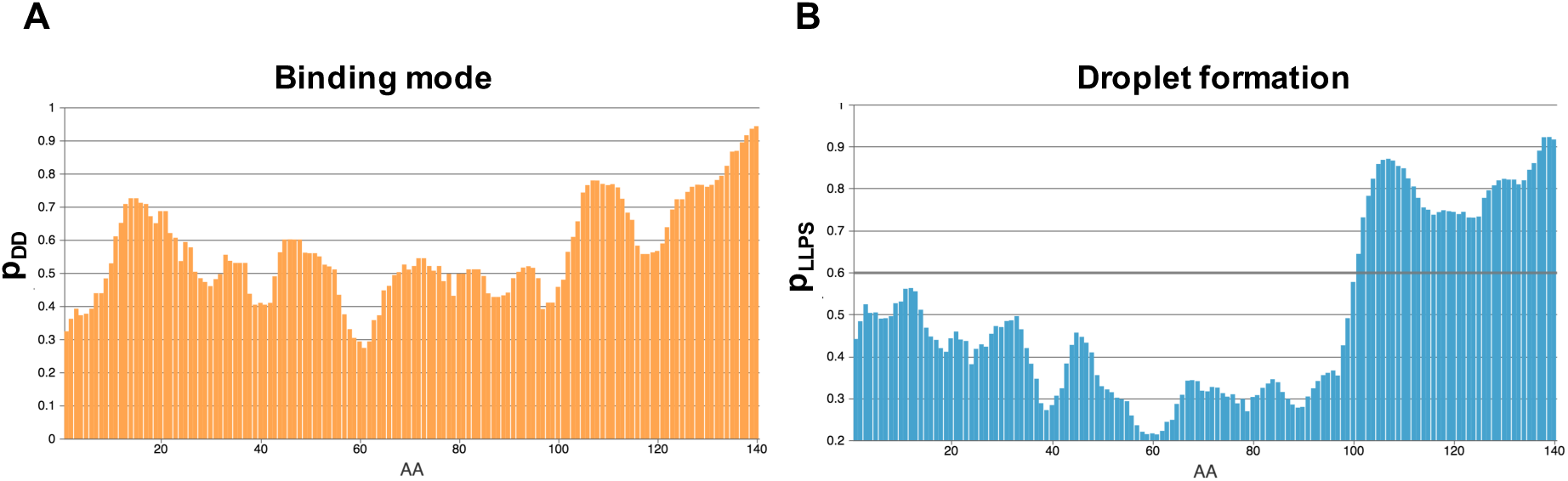
The acidic C-terminal region of α-synuclein likely drives droplet formation through disordered interactions. (**A**) A sequence-based prediction of the binding modes of disordered α-synuclein [73] suggests that the C-terminus remains predominantly disordered (p_DD_, orange) in complexes. (**B**) A sequence-based prediction of the probability of spontaneous liquid-liquid phase separation (p_LLPS_) indicates an important role for the C-terminus of α-synuclein in droplet formation (protdyn-fuzpred.org).

**Movie S1.** 1,6-hexanediol dissolves non-amyloid α-synuclein inclusions in *C. elegans.* Control (*left*) represents untreated animal. Hexanediol (*right*) represents animal treated with 10% (w/v) 1,6 hexanediol.

**Movie S2.** Rapid droplet formation of α-synuclein (100 μM) in the presence of 10% PEG after deposition on a glass slide starting at the edge of the solution. The scale bar represents 5 μm.

**Movie S3.** Fusion event of two α-synuclein droplets after 5 min incubation. The scale bar represents 1 μm.

**Movie S4.** Fluorescence recovery of α-synuclein after photobleaching of a region within a droplet.

**Movie S5**. Droplet formation of α-synuclein (100 μM) incubated with DOPE:DOPC:DOPS liposomes (1 mol% DOPE-Cy5) in the presence of 10% PEG, starting at the edge of the solution and spreading inwards. Droplets sediment on the glass surface over time. Channels represent α-synuclein (*left*), liposomes (*middle*) and an overlay (*right*). Scale bar represents 5 μm.

**Movie S6.** 3D rendering of a z-stack of a single α-synuclein/lipid droplet. Channels represent an overlay (*left*), α-synuclein (*middle*) and lipids (*right*). Scale bar represents 1 μm.

## References

1. Dawson TM, Dawson VL. Molecular pathways of neurodegeneration in Parkinson’s disease. Science. 2003;302(5646):819–22.

2. Poewe W, Seppi K, Tanner CM, Halliday GM, Brundin P, Volkmann J, et al. Parkinson disease. Nat Rev Dis Primers. 2017;3(1):1–21.

3. Polymeropoulos MH, Lavedan C, Leroy E, Ide SE, Dehejia A, Dutra A, et al. Mutation in the α-synuclein gene identified in families with Parkinson’s disease. Science. 1997;276(5321):2045–7.

4. Singleton A, Farrer M, Johnson J, Singleton A, Hague S, Kachergus J, et al. α-Synuclein locus triplication causes Parkinson’s disease. Science. 2003;302(5646):841-.

5. Fusco G, Pape T, Stephens AD, Mahou P, Costa AR, Kaminski CF, et al. Structural basis of synaptic vesicle assembly promoted by α-synuclein. Nat Comm. 2016;7(1):1–12.

6. Bendor JT, Logan TP, Edwards RH. The function of α-synuclein. Neuron. 2013;79(6):1044–66.

7. Burré J, Sharma M, Tsetsenis T, Buchman V, Etherton MR, Südhof TC. α-Synuclein promotes SNARE-complex assembly in vivo and in vitro. Science. 2010;329(5999):1663–7.

8. Lautenschläger J, Stephens AD, Fusco G, Ströhl F, Curry N, Zacharopoulou M, et al. C-terminal calcium binding of α-synuclein modulates synaptic vesicle interaction. Nat Comm. 2018;9(1):1–13.

9. Dehay B, Bourdenx M, Gorry P, Przedborski S, Vila M, Hunot S, et al. Targeting α-synuclein for treatment of Parkinson’s disease: mechanistic and therapeutic considerations. Lancet Neurol. 2015;14(8):855–66.

10. Brundin P, Dave KD, Kordower JH. Therapeutic approaches to target alpha-synuclein pathology. Exp Neurol. 2017;298:225–35.

11. Valera E, Masliah E. Therapeutic approaches in Parkinson’s disease and related disorders. J Neurochem. 2016;139:346–52.

12. Schapira AH, Olanow CW, Greenamyre JT, Bezard E. Slowing of neurodegeneration in Parkinson’s disease and Huntington’s disease: future therapeutic perspectives. Lancet. 2014;384(9942):545–55.

13. Goedert M, Jakes R, Spillantini MG. The synucleinopathies: twenty years on. J Parkinson’s Dis. 2017;7(s1):S51–S69.

14. Spillantini MG, Schmidt ML, Lee VM-Y, Trojanowski JQ, Jakes R, Goedert M. α-Synuclein in Lewy bodies. Nature. 1997;388(6645):839–40.

15. Spillantini MG, Crowther RA, Jakes R, Hasegawa M, Goedert M. α-Synuclein in filamentous inclusions of Lewy bodies from Parkinson’s disease and dementia with Lewy bodies. Proc Natl Acad Sci USA. 1998;95(11):6469–73.

16. Mahul-Mellier A-L, Burtscher J, Maharjan N, Weerens L, Croisier M, Kuttler F, et al. The process of Lewy body formation, rather than simply α-synuclein fibrillization, is one of the major drivers of neurodegeneration. Proc Natl Acad Sci USA. 2020;117(9):4971–82.

17. Shahmoradian SH, Lewis AJ, Genoud C, Hench J, Moors TE, Navarro PP, et al. Lewy pathology in Parkinson’s disease consists of crowded organelles and lipid membranes. Nat Neurosci. 2019;22(7):1099–109.

18. Wakabayashi K, Tanji K, Odagiri S, Miki Y, Mori F, Takahashi H. The Lewy Body in Parkinson’s Disease and Related Neurodegenerative Disorders. Molecular Neurobiology. 2013;47(2):495–508. doi: 10.1007/s12035-012-8280-y.

19. Arima K, Uéda K, Sunohara N, Hirai S, Izumiyama Y, Tonozuka-Uehara H, et al. Immunoelectron-microscopic demonstration of NACP/α-synuclein-epitopes on the filamentous component of Lewy bodies in Parkinson’s disease and in dementia with Lewy bodies. Brain Research. 1998;808(1):93–100. doi: 10.1016/s0006-8993(98)00734-3.

20. Spillantini MG, Crowther RA, Jakes R, Hasegawa M, Goedert M. -Synuclein in filamentous inclusions of Lewy bodies from Parkinson’s disease and dementia with Lewy bodies. Proceedings of the National Academy of Sciences. 1998;95(11):6469–73. doi: 10.1073/pnas.95.11.6469.

21. Davis AA. Beyond synuclein: Organelle accumulation in Lewy bodies may drive neurodegeneration. Sci Transl Med. 2020;12(536):eabb2778.

22. Li B, Ge P, Murray KA, Sheth P, Zhang M, Nair G, et al. Cryo-EM of full-length α-synuclein reveals fibril polymorphs with a common structural kernel. Nat Comm. 2018;9(1):1–10.

23. Cremades N, Cohen SI, Deas E, Abramov AY, Chen AY, Orte A, et al. Direct observation of the interconversion of normal and toxic forms of α-synuclein. Cell. 2012;149(5):1048–59.

24. Iljina M, Garcia GA, Horrocks MH, Tosatto L, Choi ML, Ganzinger KA, et al. Kinetic model of the aggregation of alpha-synuclein provides insights into prion-like spreading. Proceedings of the National Academy of Sciences. 2016;113(9):E1206–E15. doi: 10.1073/pnas.1524128113.

25. Cohen SIA, Vendruscolo M, Dobson CM, Knowles TPJ. Nucleated polymerization with secondary pathways. II. Determination of self-consistent solutions to growth processes described by non-linear master equations. The Journal of Chemical Physics. 2011;135(6):065106. doi: 10.1063/1.3608917.

26. Knowles TPJ, Waudby CA, Devlin GL, Cohen SIA, Aguzzi A, Vendruscolo M, et al. An Analytical Solution to the Kinetics of Breakable Filament Assembly. Science. 2009;326(5959):1533–7. doi: 10.1126/science.1178250.

27. Schweighauser M, Shi Y, Tarutani A, Kametani F, Murzin AG, Ghetti B, et al. Structures of α-synuclein filaments from multiple system atrophy. BioRxiv. 2020.

28. Van Ham TJ, Thijssen KL, Breitling R, Hofstra RMW, Plasterk RHA, Nollen EAA. C. elegans Model Identifies Genetic Modifiers of α-Synuclein Inclusion Formation During Aging. PLoS Genetics. 2008;4(3):e1000027. doi: 10.1371/journal.pgen.1000027.

29. Visanji NP, Brotchie JM, Kalia LV, Koprich JB, Tandon A, Watts JC, et al. α-Synuclein-Based Animal Models of Parkinson’s Disease: Challenges and Opportunities in a New Era. Trends Neurosci. 2016;39(11):750-62. Epub 2016/10/26. doi: 10.1016/j.tins.2016.09.003. PubMed PMID: 27776749.

30. Volpicelli-Daley LA, Luk KC, Lee VM. Addition of exogenous α-synuclein preformed fibrils to primary neuronal cultures to seed recruitment of endogenous α-synuclein to Lewy body and Lewy neurite-like aggregates. Nat Protoc. 2014;9(9):2135-46. Epub 2014/08/15. doi: 10.1038/nprot.2014.143. PubMed PMID: 25122523; PubMed Central PMCID: PMCPMC4372899.

31. Ko LW, Ko HH, Lin WL, Kulathingal JG, Yen SH. Aggregates assembled from overexpression of wild-type alpha-synuclein are not toxic to human neuronal cells. J Neuropathol Exp Neurol. 2008;67(11):1084-96. Epub 2008/10/30. doi: 10.1097/NEN.0b013e31818c3618. PubMed PMID: 18957893; PubMed Central PMCID: PMCPMC2768257.

32. Taschenberger G, Garrido M, Tereshchenko Y, Bähr M, Zweckstetter M, Kügler S. Aggregation of αSynuclein promotes progressive in vivo neurotoxicity in adult rat dopaminergic neurons. Acta Neuropathol. 2012;123(5):671-83. Epub 2011/12/15. doi: 10.1007/s00401-011-0926-8. PubMed PMID: 22167382; PubMed Central PMCID: PMCPMC3316935.

33. Hyman AA, Weber CA, Jülicher F. Liquid-liquid phase separation in biology. Annu Rev Cell Dev Biol. 2014;30:39–58.

34. Banani SF, Lee HO, Hyman AA, Rosen MK. Biomolecular condensates: organizers of cellular biochemistry. Nat Rev Mol Cell Biol. 2017;18(5):285–98.

35. Shin Y, Brangwynne CP. Liquid phase condensation in cell physiology and disease. Science. 2017;357(6357):eaaf4382.

36. Boeynaems S, Alberti S, Fawzi NL, Mittag T, Polymenidou M, Rousseau F, et al. Protein phase separation: a new phase in cell biology. Trends Cell Biol. 2018;28(6):420–35.

37. Ader C, Frey S, Maas W, Schmidt HB, Görlich D, Baldus M. Amyloid-like interactions within nucleoporin FG hydrogels. Proc Natl Acad Sci USA. 2010;107(14):6281–5.

38. Patel A, Lee HO, Jawerth L, Maharana S, Jahnel M, Hein MY, et al. A liquid-to-solid phase transition of the ALS protein FUS accelerated by disease mutation. Cell. 2015;162(5):1066–77.

39. Murakami T, Qamar S, Lin JQ, Schierle GSK, Rees E, Miyashita A, et al. ALS/FTD mutation-induced phase transition of FUS liquid droplets and reversible hydrogels into irreversible hydrogels impairs RNP granule function. Neuron. 2015;88(4):678–90.

40. Ambadipudi S, Biernat J, Riedel D, Mandelkow E, Zweckstetter M. Liquid–liquid phase separation of the microtubule-binding repeats of the Alzheimer-related protein Tau. Nat Comm. 2017;8(1):1–13.

41. Wegmann S, Eftekharzadeh B, Tepper K, Zoltowska KM, Bennett RE, Dujardin S, et al. Tau protein liquid–liquid phase separation can initiate tau aggregation. EMBO J. 2018;37(7).

42. Kanaan NM, Hamel C, Grabinski T, Combs B. Liquid-liquid phase separation induces pathogenic tau conformations in vitro. Nature Communications. 2020;11(1). doi: 10.1038/s41467-020-16580-3.

43. Ray S, Singh N, Kumar R, Patel K, Pandey S, Datta D, et al. α-Synuclein aggregation nucleates through liquid–liquid phase separation. Nat Chem. 2020. doi: 10.1038/s41557-020-0465-9.

44. Laine RF, Sinnige T, Ma KY, Haack AJ, Poudel C, Gaida P, et al. Fast fluorescence lifetime imaging reveals the aggregation processes of α-synuclein and polyglutamine in aging Caenorhabditis elegans. ACS Chem Biol. 2019;14(7):1628–36.

45. Poudel C, Mela I, Kaminski CF. High-throughput, multi-parametric, and correlative fluorescence lifetime imaging. Methods Appl Fluoresc. 2020;8(2):024005.

46. Kaminski Schierle GS, Bertoncini CW, Chan FT, van der Goot AT, Schwedler S, Skepper J, et al. A FRET sensor for non-invasive imaging of amyloid formation in vivo. ChemPhysChem. 2011;12(3):673–80.

47. Van Ham TJ, Thijssen KL, Breitling R, Hofstra RM, Plasterk RH, Nollen EA. C. elegans model identifies genetic modifiers of α-synuclein inclusion formation during aging. PLoS Gen. 2008;4(3).

48. Kroschwald S, Maharana S, Simon A. Hexanediol: a chemical probe to investigate the material properties of membrane-less compartments. Matters. 2017;3(5):e201702000010.

49. Molliex A, Temirov J, Lee J, Coughlin M, Kanagaraj AP, Kim HJ, et al. Phase separation by low complexity domains promotes stress granule assembly and drives pathological fibrillization. Cell. 2015;163(1):123–33.

50. Harmon TS, Holehouse AS, Rosen MK, Pappu RV. Intrinsically disordered linkers determine the interplay between phase separation and gelation in multivalent proteins. Elife. 2017;6:e30294.

51. Burré J, Sharma M, Südhof TC. α-Synuclein assembles into higher-order multimers upon membrane binding to promote SNARE complex formation. Proc Natl Acad Sci USA. 2014;111(40):E4274–E83.

52. Takamori S, Holt M, Stenius K, Lemke EA, Grønborg M, Riedel D, et al. Molecular anatomy of a trafficking organelle. Cell. 2006;127(4):831–46.

53. Galvagnion C, Brown JW, Ouberai MM, Flagmeier P, Vendruscolo M, Buell AK, et al. Chemical properties of lipids strongly affect the kinetics of the membrane-induced aggregation of α-synuclein. Proc Natl Acad Sci USA. 2016;113(26):7065–70.

54. Chen X, Wu X, Wu H, Zhang M. Phase separation at the synapse. Nat Neurosci. 2020:1–10.

55. Milovanovic D, Wu Y, Bian X, De Camilli P. A liquid phase of synapsin and lipid vesicles. Science. 2018;361(6402):604–7.

56. Chen X, Wu X, Wu H, Zhang M. Phase separation at the synapse. Nature Neuroscience. 2020. doi: 10.1038/s41593-019-0579-9.

57. Atias M, Tevet Y, Sun J, Stavsky A, Tal S, Kahn J, et al. Synapsins regulate α-synuclein functions. Proc Natl Acad Sci USA. 2019;116(23):11116–8.

58. Zaltieri M, Grigoletto J, Longhena F, Navarria L, Favero G, Castrezzati S, et al. α-synuclein and synapsin III cooperatively regulate synaptic function in dopamine neurons. J Cell Sci. 2015;128(13):2231–43.

59. Longhena F, Faustini G, Varanita T, Zaltieri M, Porrini V, Tessari I, et al. Synapsin III is a key component of α-synuclein fibrils in Lewy bodies of PD brains. Brain Pathology. 2018;28(6):875–88.

60. Araki K, Yagi N, Aoyama K, Choong C-J, Hayakawa H, Fujimura H, et al. Parkinson’s disease is a type of amyloidosis featuring accumulation of amyloid fibrils of α-synuclein. Proc Natl Acad Sci USA. 2019;116(36):17963–9.

61. Boke E, Ruer M, Wühr M, Coughlin M, Lemaitre R, Gygi SP, et al. Amyloid-like self-assembly of a cellular compartment. Cell. 2016;166(3):637–50.

62. Yuan C, Levin A, Chen W, Xing R, Zou Q, Herling TW, et al. Nucleation and Growth of Amino Acid and Peptide Supramolecular Polymers through Liquid–Liquid Phase Separation. Angew Chem Int Ed. 2019;131(50):18284–91.

63. Pogostin BH, Linse S, Olsson U. Fibril Charge Affects α-Synuclein Hydrogel Rheological Properties. Langmuir. 2019;35(50):16536–44.

64. Kumar R, Das S, Mohite GM, Rout SK, Halder S, Jha NN, et al. Cytotoxic Oligomers and Fibrils Trapped in a Gel-like State of α-Synuclein Assemblies. Angew Chem Int Ed. 2018;57(19):5262–6.

65. Buell AK, Galvagnion C, Gaspar R, Sparr E, Vendruscolo M, Knowles TP, et al. Solution conditions determine the relative importance of nucleation and growth processes in α-synuclein aggregation. Proc Natl Acad Sci USA. 2014;111(21):7671–6.

66. Galvagnion C, Buell AK, Meisl G, Michaels TC, Vendruscolo M, Knowles TP, et al. Lipid vesicles trigger α-synuclein aggregation by stimulating primary nucleation. Nat Chem Biol. 2015;11(3):229.

67. Nguyen M, Krainc D. LRRK2 phosphorylation of auxilin mediates synaptic defects in dopaminergic neurons from patients with Parkinson’s disease. Proceedings of the National Academy of Sciences. 2018;115(21):5576–81. doi: 10.1073/pnas.1717590115.

68. Goldstein B. Sydney Brenner on the Genetics of Caenorhabditis elegans. Genetics. 2016;204(1):1.

69. Chow C, Kaneko T. Automatic boundary detection of the left ventricle from cineangiograms. Comput Biomed Res. 1972;5(4):388–410.

70. Warren SC, Margineanu A, Alibhai D, Kelly DJ, Talbot C, Alexandrov Y, et al. Rapid global fitting of large fluorescence lifetime imaging microscopy datasets. PLoS One. 2013;8(8).

71. Bhaskaran S, Butler JA, Becerra S, Fassio V, Girotti M, Rea SL. Breaking Caenorhabditis elegans the easy way using the Balch homogenizer: an old tool for a new application. Anal Biochem. 2011;413(2):123–32.

72. Hoyer W, Antony T, Cherny D, Heim G, Jovin TM, Subramaniam V. Dependence of α-synuclein aggregate morphology on solution conditions. J Mol Biol. 2002;322(2):383–93.

73. Miskei M, Horváth A, Vendruscolo M, Fuxreiter M. Sequence-based determinants and prediction of fuzzy interactions in protein complexes. J Mol Biol. 2020.

